# Eip74EF is a dominant modifier for ALS-FTD linked mutant VCP phenotypes in *Drosophila*, but not miR-34

**DOI:** 10.1101/2020.11.20.375360

**Authors:** Madeleine R. Chalmers, JiHye Kim, Nam Chul Kim

**Affiliations:** Department of Pharmacy Practice and Pharmaceutical Sciences, College of Pharmacy, University of Minnesota, Duluth, MN 55812

## Abstract

In 2012 Liu *et al*. reported that miR-34 is an age-related miRNA regulating age-associated events and long-term brain integrity in *Drosophila*. They demonstrated that modulating miR-34 and its downstream target Eip74EF showed beneficial effects on age-related diseases using a *Drosophila* model of SCA3trQ78. These results imply that miR-34 could be a general genetic modifier and therapeutic candidate for age-related diseases. Therefore, we examined the effect of miR-34 and Eip74EF on another age-related *Drosophila* disease model. Using a *Drosophila* model expressing mutant *Drosophila* VCP that causes amyotrophic lateral sclerosis (ALS) and frontotemporal dementia (FTD), or multisystem proteinopathy (MSP), we demonstrated that abnormal eye phenotypes generated by *Drosophila* VCP R152H were rescued when expressed with Eip74EF siRNA. Contrary to our expectation, miR-34 overexpression resulted in lethality when expressed with mutant VCP. Our data indicate that the other downstream targets of miR-34 might more significantly interact with mutant VCP, causing lethality. Identifying transcriptional targets of Eip74EF might provide valuable insights into diseases caused by mutations in VCP such as ALS, FTD, and MSP.

## Introduction

Multisystem proteinopathy (MSP) is a devastating age-related familial disease with no effective treatments. MSP is a dominantly inherited autosomal disease with differing degrees of severity, ranging in different combinations of inclusion body myopathy (IBM), Paget’s disease, frontotemporal dementia (FTD), and amyotrophic lateral sclerosis (ALS) in the same family members (Nalbandian *et al*. 2011; Benatar *et al*. 2013). Thus, it has been known as IBMPFD-ALS. Around 70% of MSP patients have a missense mutation in the gene encoding for valosin-containing protein (VCP) (Takada 2015). VCP is a ubiquitin-dependent segregase with a wide range of functions, some of which include protein degradation, nuclear envelope construction, and assembly of the Golgi apparatus and endoplasmic reticulum (Ritson *et al*. 2010). The age of onset for MSP patients ranges from late thirties to mid-fifties (Benatar *et al*. 2013; Al-Obeidi *et al*. 2018).

Despite no readily available treatments, regulatory RNAs might be a potential therapeutic technique for degenerative diseases such as MSP (Campbell and Booth 2015; Salta and De Strooper 2017). MicroRNAs (miRNA) are notable as they have a wide degree of functions and are essential for development, including cell differentiation and aging (Liu *et al*. 2012; Gebert and MacRae 2019). Interestingly, Liu *et al*. identified that microRNA-34 (miR-34) is involved in the aging process in *Drosophila melanogaster* by a miRNA microarray analysis. When miR-34 was knocked out in *Drosophila*, life spans were reduced, and aging deficits were present including accelerated neurodegeneration. When miR-34 was overexpressed in *Drosophila* the lifespan of adult *Drosophila* increased. Therefore, miR-34 can potentially protect against neurodegeneration and extend the lifespan. Indeed, they tested miR-34 against *Drosophila* expressing the polyQ domain of mutant Ataxin-3 (SCA3trQ78) causing spinocerebellar ataxia, and found that inclusion formation was slowed and neurodegeneration was suppressed. Finally, they identified Eip74EF as a downstream target of miR-34 to show that miR-34 has protective effects against aging and neurodegeneration (Liu *et al*. 2012). Ecdysone-induced protein 74EF (Eip74EF) encodes a transcription factor, E74A, involved in ecdysone-mediated cell death (Liu *et al*. 2012; Nicolson *et al*. 2015). E74A is essential for development but is harmful when present in adult *Drosophila*. Liu *et al*. demonstrated that miR-34 targets Eip74EF and decreases E74A levels which resulted in protective effects against aging and neurodegeneration, while also increasing the median lifespan of adult *Drosophila*. Their findings provide an intriguing possibility that miR-34 or siRNAs against Eip74EF may have roles in age-related degenerative diseases such as MSP. Thus, we hypothesized that miR-34 and its downstream target, Eip74EF, are genetic modifiers for mutant VCP causing age-related MSP. Importantly, the evolutionarily conserved miR-34 might also act as a genetic modifier in humans and may be a potential candidate for therapeutics for mutant VCP-induced diseases.

## Results and Discussion

To test this hypothesis, we examined genetic interactions between mutant VCP and miR-34 pathways. We used a previously published *Drosophila* model expressing mutant *Drosophila* VCP R152H (dVCP R152H) in eyes with GMR-GAL4 (Ritson *et al*. 2010). This model has been successfully used to find genetic modifiers modulating mutant VCP pathogenesis (Ritson *et al*. 2010; Kim *et al*. 2013). When dVCP R152H was expressed in *Drosophila* eyes with a GMR-GAL4 driver, severe eye degeneration was observed as published (Ritson *et al*. 2010). We examined the effect of miR-34 overexpression and RNAi-mediated knockdown for Eip74EF, using this rough eye phenotype. Based on the findings of Liu *et al*., we expected to see a rescue in the degenerative eye phenotype when Eip74EF had reduced expression, as well as a rescue in the eye phenotype when miR-34 was overexpressed. *Drosophila* expressing dVCP R152H in their eyes (GMR-GAL4, UAS dVCP R152H/CyO) were crossed with *Drosophila* lines carrying a construct for RNAi of Eip74EF (Bloomington 29353) and miR-34 overexpression (Bloomington 63790, Bloomington 41158), respectively. mCherry RNAi control line (Bloomington 35785) was used as a negative control. Eip74EF siRNA expression rescued abnormal eye phenotypes generated by dVCP R152H compared to the degenerated eyes of control siRNA expressing (mCherry siRNA) *Drosophila* (Figure 1a). However, both lines of miR-34 overexpression *Drosophila* resulted in lethality (Figure 1a). Male Eip74EF siRNA progeny exhibited significantly rescued eye phenotypes compared to male mCherry siRNA progeny (P = 0.0008), while female Eip74EF siRNA progeny also exhibited significantly rescued eye phenotypes compared to female mCherry siRNA progeny (P = 0.003)(Figure 1b). Thus, contrary to our expectation, we did not observe a rescue in the degenerative eye phenotype of miR-34 overexpressing progeny, however, we did observe a rescue in the degenerative eye phenotype of Eip74EF siRNA progeny.

**Figure 1.**
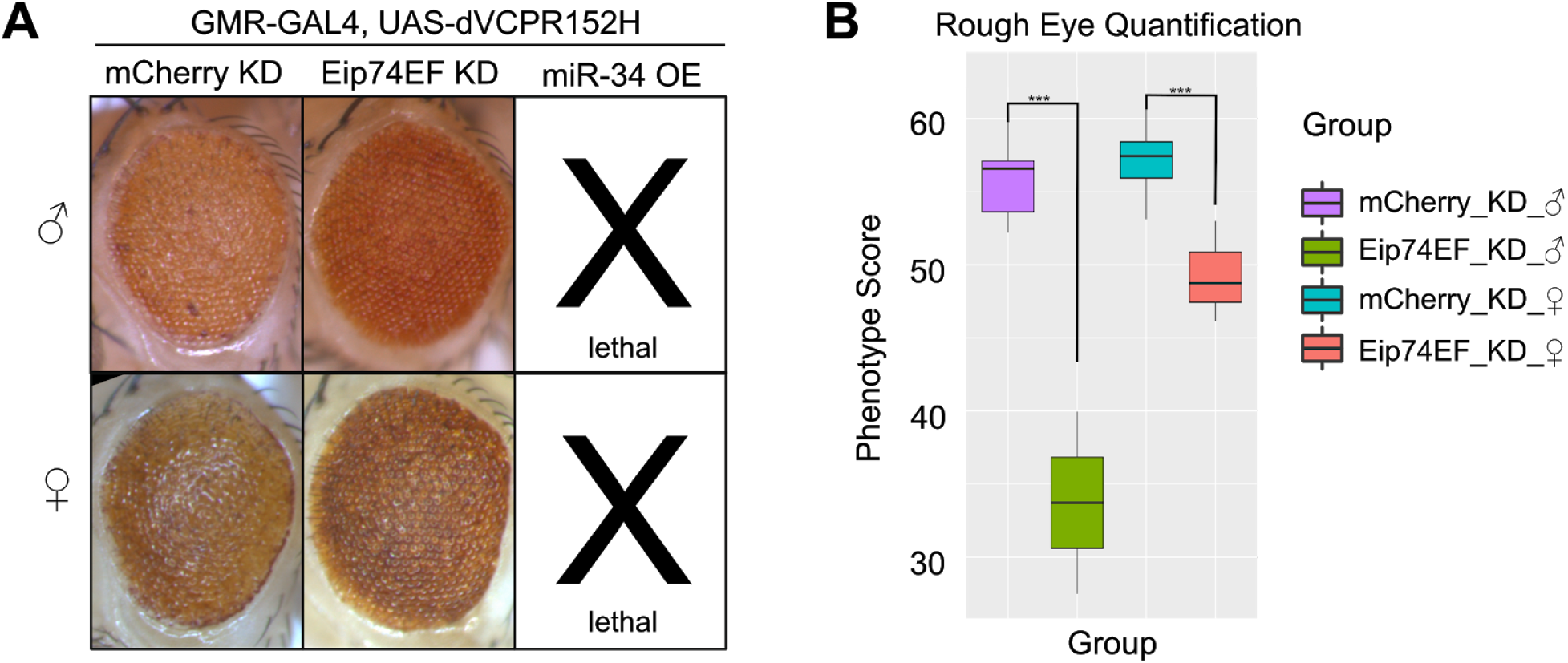
Eip74EF siRNA is a genetic modifier for mutant VCP and significantly rescues degenerative phenotypes while miR-34 overexpression resulted in lethality: **A.** Eip74EF siRNA (KD) expression significantly reduced mutant dVCP-induced abnormal eye phenotypes compared to a control mCherry siRNA (KD) expression. miR-34 overexpression (OE) with dVCP R152H resulted in complete lethality. **B.** Severity score quantification of *Drosophila* eyes. The effect of Eip74EF siRNA expression on dVCP R152H is statistically significant in both male and female progeny (***: P = 0.0008 and P = 0.0003, respectively).

There can be some possible explanations for this discrepancy. We used UAS-miR-34 lines from the Bloomington Stock Center, not the same miR-34 lines created by Liu *et al* (2012). These lines likely have different expression levels of miR-34 when it is expressed with GMR-GAL4.

The higher expression of miR-34 may result in pupal lethality in progenies carrying dVCP R152H and miR-34 together. However, please note that RNAi-mediated knockdown of Eip74EF successfully rescued dVCP R152H-mediated toxicity though we did not use a hypomorphic mutant used by Liu *et al*. Thus, the higher expression of miR-34 may induce lethality via other genes regulated by miR-34, not Eip74EF. The toxicity of dVCP R152H expressed by GMR-GAL4 is high from the late pupal stage to the early adult stage which is regulated by the ecdysone-mediated transcriptional program. Therefore, it is not surprising that the reduction of Eip74EF, an ecdysone-activated transcription factor, involved in apoptosis, ameliorates dVCP R152H toxicity. However, in the case of miR-34 with dVCP R152H, combining overexpression of dVCP R152H and miR-34 in this stage may cause further complex genetic interactions. miR-34 might induce lethality through its downstream targets, suppressing apoptosis, such as SIRT1 (Yamakuchi and Lowenstein 2009), while still suppressing Eip74EF expression that is beneficial for dVCP R152H-mediated toxicity. If we use a *Drosophila* model expressing dVCP R152H in the late adult stage, we might observe the expected beneficial effects of miR-34 overexpression.

Together, our data suggests that Eip74EF, not miR-34, is a strong genetic modifier for mutant VCP that causes age-related degenerative diseases including ALS, FTD and MSP. Although Eip74EF is not well conserved in humans, further studies identifying downstream transcriptional networks activated by Eip74EF might provide valuable insights in what cellular pathways are involved in mutant VCP-caused diseases and how to treat these diseases.

## Methods

*Drosophila* were crossed and grown on Bloomington Stock Center standard cornmeal food at 25°C. Progeny were collected at two days old, and *Drosophila* eye pictures were taken using a Leica M205C stereomicroscope with a Leica DFC320 digital camera. Eye quantification was performed using softwares ilastik (Berg *et al*. 2019) and Flynotyper (Iyer *et al*. 2016). A Student’s t-test was used to compare the phenotypic severity of mCherry siRNA and Eip74EF siRNA *Drosophila*.

### *Drosophila* lines

All lines were obtained from the Bloomington Drosophila Stock Center.

35785 y[1] sc[*] v[1] sev[21]; P{y[+t7.7] v[+t1.8]=VALIUM20-mCherry}attP2: expresses siRNA for mCherry under UAS control in the VALIUM20 vector.

29353 y[1] v[1]; P{y[+t7.7] v[+t1.8]=TRiP.JF02515}attP2: expresses siRNA for Eip74EF (FBgn0000567) under UAS control in the VALIUM10 vector.

63790 w[*]; P{w[+mC]=UAS-DsRed-mir-34.L}3/TM3, Sb[1] Ser[1]: expresses mir-34 under UAS control.

41158 w[1118]; P{y[+t7.7] w[+mC]=UAS-LUC-mir-34.T}attP2: expresses mir-34 (FBgn0262459) under UAS control.

*Drosophila* line GMR-GAL4, UAS dVCP R152H/CyO, was a gift from Dr. J. Paul Taylor.

## Acknowledgments

We thank the Bloomington Drosophila Stock Center for *Drosophila* stocks. We also thank Minwoo Baek and Swati Maitra for their feedback during the writing process.

## Funding

Financial support was provided by a Muscular Dystrophy Association grant to Nam Chul Kim

## Notes

### Competing Interest Statement

The authors have declared no competing interest.

